# Computational characterization of recombinase circuits for periodic behaviors

**DOI:** 10.1101/2021.11.06.467548

**Authors:** Judith Landau, Christian Cuba Samaniego, Giulia Giordano, Elisa Franco

## Abstract

In nature, recombinases are site-specific proteins capable of rearranging DNA, and they are expanding the repertoire of gene editing tools used in synthetic biology. The on/off response of recombinases, achieved by inverting the direction of a promoter, makes them suitable for Boolean logic computation; however, recombinase-based logic gate circuits are single-use due to the irreversibility of the DNA rearrangement, and it is still unclear how a dynamical circuit, such as an oscillator, could be engineered using recombinases. Preliminary work has demonstrated that recombinase-based circuits can yield periodic behaviors in a deterministic setting. However, since a few molecules of recombinase are enough to perform the inverting function, it is crucial to assess how the inherent stochasticity at low copy number affects the periodic behavior. Here, we propose six different circuit designs for recombinase-based oscillators. We model them in a stochastic setting, leveraging the Gillespie algorithm for extensive simulations, and we show that they can yield periodic behaviors. To evaluate the incoherence of oscillations, we use a metric based on the statistical properties of auto-correlation functions. The main core of our design consists of two self-inhibitory, recombinase-based modules coupled by a common promoter. Since each recombinase inverts its own promoter, the overall circuit can give rise to switching behavior characterized by a regular period. We introduce different molecular mechanisms (transcriptional regulation, degradation, sequestration) to tighten the control of recombinase levels, which slows down the response timescale of the system and thus improves the coherence of oscillations. Our results support the experimental realization of recombinase-based oscillators and, more generally, the use of recombinases to generate dynamic behaviors in synthetic biology.

## 1 Introduction

Oscillatory behaviors drive essential processes in nature. For example, the mitotic oscillator drives cell division [1, 2, 3], the circadian oscillator drives the sleep-wake cycle [4, 5, 6], and the segmentation clock drives spatial pattern formation during vertebrate embryonic development [7, 8]. These numerous examples have motivated biologists, physicists, and mathematicians to look for the design principles required to build biomolecular oscillators from the bottom-up [9, 10], and many decades of theoretical and experimental research have consolidated design principles for oscillator design, which primarily include the presence of a negative feedback loop and of mechanisms for local destabilization, such as positive feedback and delays [11, 12, 13, 14]. The implementation of these design principles within artificial genetic circuits has been shown to facilitate the emergence of periodic behaviors, in contexts spanning from single cell metabolism to multi-cellular environments [15]. However, while tremendous progress has been made toward building robust synthetic oscillators using transcription factors [16, 17], demonstrations of new oscillator architectures are lagging. This is in contrast with the dramatic expansion of molecular functions and gene editing tools harnessed by synthetic biology [18].

Recombinases are a class of proteins with major potential toward engineering cellular behavior [19]. These enzymes cleave and rejoin DNA strands with high specificity for given genetic domains (sequences), and many orthogonal recombinases exist. By carefully placing these domains, recombinases can perform diverse operations such as DNA excision, insertion, and translocation to generate logic and regulatory circuits that are easy to scale, and can be implemented in a variety of organisms [20, 21]. Because recombinases make it possible to swap domains of DNA, they can “rewire” entire gene expression pathways. By simply inverting target promoter regions, recombinases can activate or deactivate expression with a nonlinear response that is comparable to a digital on/off switch. While this is an attractive feature toward building complex cellular circuits, the use of recombinases to generate periodic behaviors has received little attention. Creating periodic cycles of DNA site inversion using recombinases has a fundamental limitation posed by the difficulty to reverse-rearrange DNA. This limitation can be overcome by adopting serine integrases (Box 1) which allow for reversible rearrangement of DNA, as shown in recent work toward the design of toggle switches and counters [22, 23].

In this paper, we describe and compare various candidate architectures to design oscillators using recombinases that generate regulatory feedback loops with nonlinear, switch-like responses. The simplest recombinase-based oscillator could be described as the interconnection of two negative feedback loops. The design includes a single promoter between recombinase sites. When the promoter points to the right, the first recombinase is expressed and causes inversion of the promoter to the left. When the promoter points to the left, it drives expression of a second recombinase that causes inversion of the promoter back to the right. Thus, each recombinase suppresses its own production. Using a model based on ordinary differential equations, we previously found that this simple circuit can support periodic switching [24]. However, in any practical implementation, just a few copies of recombinase are sufficient to cause excision and inversion, so stochastic models are needed to computationally explore the circuit behavior. Moreover, it is unclear how the inherent stochasticity at low copy numbers affects the periodic switching behavior of a single-copy recombinase-based oscillator. Using the Gillespie Algorithm, we examine alternative designs incorporating different reactions to regulate more tightly the recombinase levels in the circuit, with the goal of improving the coherence of the periodic behavior. To evaluate the period incoherence, we introduce a metric based on the variance of the computationally measured autocorrelation function. We assess the effects of various reaction rate parameters on the oscillatory behavior using our period incoherence metric, and we use it as a means to compare the different designs. Overall, we find that periodic behavior is achievable in all designs when adopting biologically plausible reaction parameters. Our findings support the experimental implementation of a new class of recombinase-based oscillators.

## 2 Methods

### Stochastic simulations

We used the Gillespie algorithm [33], implemented in MATLAB, to generate stochastic trajectories of a set of species interacting according to a list of chemical reactions. The reaction rate constants (Table 1) associated with each reaction are converted to reaction propensities, and the algorithm simulates which reactions fire at each step of the simulation. The copy number (or concentration) increases or decreases one molecule at a time while the algorithm tracks the changes in all species over time. We study six distinct chemical reaction networks that model candidate oscillators based on recombinase interactions. Every design has a promoter that inverts back and forth under the action of a serine integrase and the same serine integrase fused to its RDF, so the chemical reactions include promoter inversion, transcription, translation, degradation. The left-pointing and right-pointing configurations of the promoter are considered different species: *S*_*L*_ and *S*_*R*_, respectively. The propensity for converting from one promoter species to the other is controlled by a function of recombinase concentration since recombinase is what physically inverts the promoter. For each design we generated 500 stochastic trajectories using a reaction volume of 1 femtoliter; initial conditions were set to zero for all species except one copy of left-pointing promoter (*S*_*L*_ = 1).

**Table 1:** Nominal simulation parameters of the controlled system.

| Parameter | Description | Value | Other studies |
| --- | --- | --- | --- |
| $a$ (/s/M) | Association rate | $2.1 \times 10^4$ | $10^4 - 10^6$ [41, 42, 43] |
| $d$ (/s) | Dissociation rate | $2.1 \times 10^2$ | $1.7 \cdot 10^{-5} - 1.7 \cdot 10^{-1}$ [43] |
| $\phi$ (/s) | RNA degradation | $12 \times 10^{-4}$ | $10^{-4} - 10^{-3}$ [44] |
| $\delta$ (/s) | Protein degradation | $3.8 \times 10^{-4}$ | $10^{-4} - 10^{-3}$ [45] |
| $\rho, \rho_1$ (/s) | Translation rate | $7.5 \times 10^{-3}$ | $3 \cdot 10^{-3} - 1.5 \cdot 10^{-2}$ [46]* |
| $\theta$ (M/s) | Transcription rate | $2.78 \times 10^{-11}$ | $2.8 \cdot 10^{-11} - 2.8 \cdot 10^{-8}$ [47, 48] |
| $K$ (nM) | Dissociation constant | 200 | $10^{-2} - 10^3$ [49, 50] |
| $r$ (1/s) | Switching rate | $1.4 \times 10^{-3}$ | $8 \cdot 10^{-5} - 1.6 \cdot 10^{-2}$ [51, 30] |
| $K_d$ (nM) | Dissociation constant | 100 | $10^{-2} - 10^3$ [49, 50] |
| $\beta$ (M/s) | Transcription rate | $2.78 \times 10^{-11}$ | $2.8 \cdot 10^{-11} - 2.8 \cdot 10^{-8}$ [47, 48] |
| $\rho_2$ (/s) | Translation rate | $1.1 \times 10^{-2}$ | $3 \cdot 10^{-3} - 1.5 \cdot 10^{-2}$ [46]* |
| $\xi$ (1/s) | Degradation rate | $2.7 \times 10^{-5}$ | $2 \cdot 10^{-2}$ [52] |
| $\gamma$ (/s/M) | Sequestration rate | $2.1 \times 10^5$ | $10^4 - 10^6$ [53, 54] |
| $\mu$ (M/s) | Transcription rate | $5.56 \times 10^{-11}$ | $2.8 \cdot 10^{-11} - 2.8 \cdot 10^{-8}$ [47, 48] |
| $\alpha$ (M/s) | Transcription rate | $2.78 \times 10^{-11}$ | $2.8 \cdot 10^{-11} - 2.8 \cdot 10^{-8}$ [47, 48] |

\*For these parameters, we estimated the translation rate of proteins with a gene size between 0.5kb to 2kb and a synthesis rate of 15 aa/s.

#### Box 1. Serine integrases and their applications in synthetic circuits

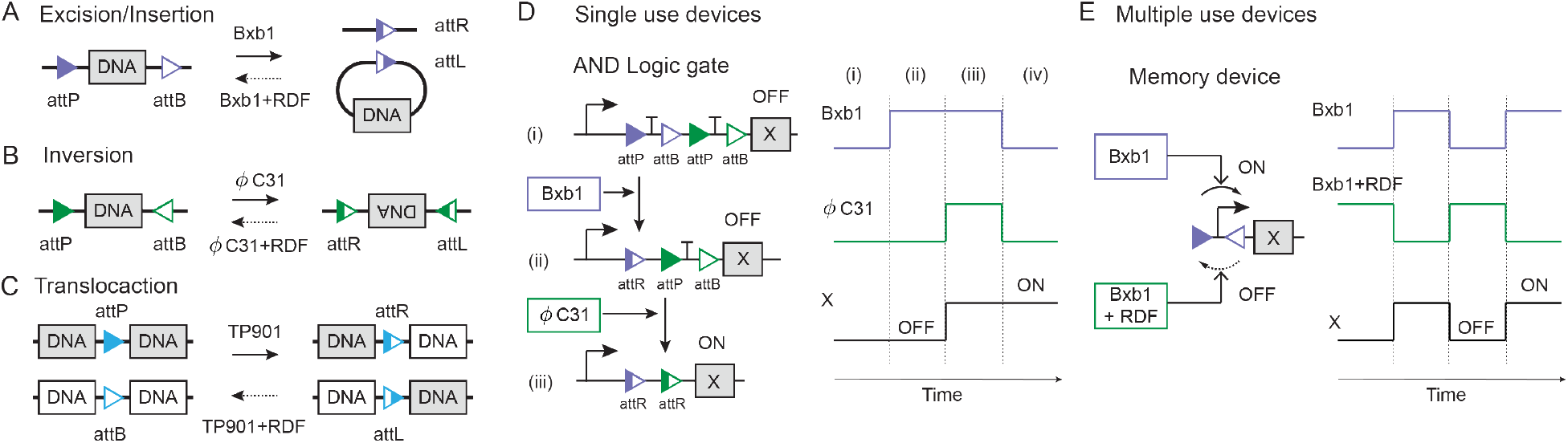

We summarize the most important functions of serine integrases (panels A-C) and their relevant applications (panels D-E). There are two families of recombinases: tyrosine recombinases and serine recombinases [19]. All recombinases are site-specific proteins that can rearrange DNA, performing, for example, excision/insertion, inversion, and translocation. Serine integrases are a subfamily of serine recombinases, each of which has a cognate Recombination Directionality Factor (RDF) that allows the serine integrase to reverse-rearrange DNA [19].

We consider three examples involving the serine integrases Bxb1, *ϕ*C31, and TP901, with cognate attP and attB binding sites. When the attP and attB binding sites have the same orientation, Bxb1 monomers form a dimer, bind to the two specific sites, and excise the DNA segment in between, as shown in panel A (excision/insertion). When the attP and attB binding sites point in opposite directions, *ϕ*C31 binds to them and inverts the DNA between the binding sites, as shown in panel B (inversion). When the attP and attB binding sites are not in the same region of DNA, TP901 can bind to the attP and attB binding sites and translocate the DNA strands, as shown in panel C (translocation). In all the above examples, adding a recombination directional factor (RDF) enables the recombinases to recognise the binding sites attL and attR present after the rearrangements described, and therefore to reverse them. Tyrosine recombinases can perform the DNA rearrangements described, among others, but not reverse them, examples including Cre, Vre, and FLP [25]. The exception to this is the pair of tyrosine recombinases FimE and HbiF that can reverse the DNA recombination completed by the other.

Recombinases have been used within logic gate circuits, such as the AND gate shown in panel D [26]. In this example, two transcription terminators are between the attP and attB binding sites for two different and orthogonal recombinases, Bxb1 and *ϕ*C31, shown in purple and green, respectively. In the absence of both recombinase inputs, the transcription of gene X is suppressed (OFF). When Bxb1 is added, it excises the first terminator. However, the transcription remains suppressed (OFF), because the second terminator is still present. When *ϕ*C31 is also added, it excises the second terminal, which finally activates the transcription of gene X (ON). Hence, two recombinase inputs (Bxb1 and *ϕ*C31) are needed to activate the circuit. One major disadvantage of recombinase-based logic gates is that the irreversibility of the DNA rearrangement means they can only be operated a single time. This circuit would require the cognate RDFs of these integrases to insert the DNA that was excised and thus be a multiple-use device.

There are few demonstrations of multiple-use, dynamic devices built using recombinases. The first example was the engineering of a programmable switch, as shown in panel E [27]. This pioneering work uses Bxb1 and its RDF to change the direction of the promoter controlling the production of X over multiple cell generations [27]. An improved version of the programmable switch uses tyrosine recombinases FimE and HbiF [28], which are the only special cases of tyrosine recombinases that allow reversible DNA rearrangement. Other dynamical circuit designs based on recombinases include a negative feedback controller to track a reference [29, 30], some theoretical designs of toggle switches that incorporate multiple copies of the circuit [22], and a single-input counting circuit [23]. The oscillatory behavior of a recombinase-based circuit has also been analyzed deterministically [24].

Because recombinases can be used as a switch to turn on/off gene expression, they are well-suited to build large Boolean logic circuits [31] that can be hierarchically composed with a predictable response [21]. In addition, self-excision recombinases were used to generate temporal responses such a pulses, and a cascade of self-excision mechanism can create a sequential pulse behavior that operates once [20]. Because leaky expression of recombinases can jeopardize circuit operation (only few protein copies are necessary to carry out their function), methods to tightly control their production are necessary, for example via light-induction [32].

### Metrics for coherence/incoherence

To evaluate the consistency/inconsistency of the period of stochastic trajectories we used an incoherence metric based on the autocorrelation function, following the approach introduced by [34, 35]. We focus strictly on the recombinase copy number and study the coherence of the oscillation period produced by each circuit design. Fully coherent oscillations not only have a regular period, but each cycle of the oscillations has comparable amplitude. In a stochastic context these requirements for perfect coherence are not achievable, so we only consider fluctuations of the period from the start of the simulation. Given a stochastic trajectory like the one in Fig. 1A, we first compute the autocorrelation function (shown in Fig. 1B) of the simulated concentration trajectory. Then we evaluate two kinds of features of the autocorrelation function: the time interval between the first peak and each consecutive peak, termed *T*_*i*_, *i* = 1, 2, …; and the time intervals between subsequent peaks, defined as Δ*T*_*i*_ = *T*_*i*_ – *T*_*i*–1_ for *i* ≥ 2, and Δ*T*_1_ = *T*_1_. As an example, the intervals Δ*T*_*i*_ and *T*_*i*_ are marked in Fig. 1B for the first three peaks.

**Figure 1:**
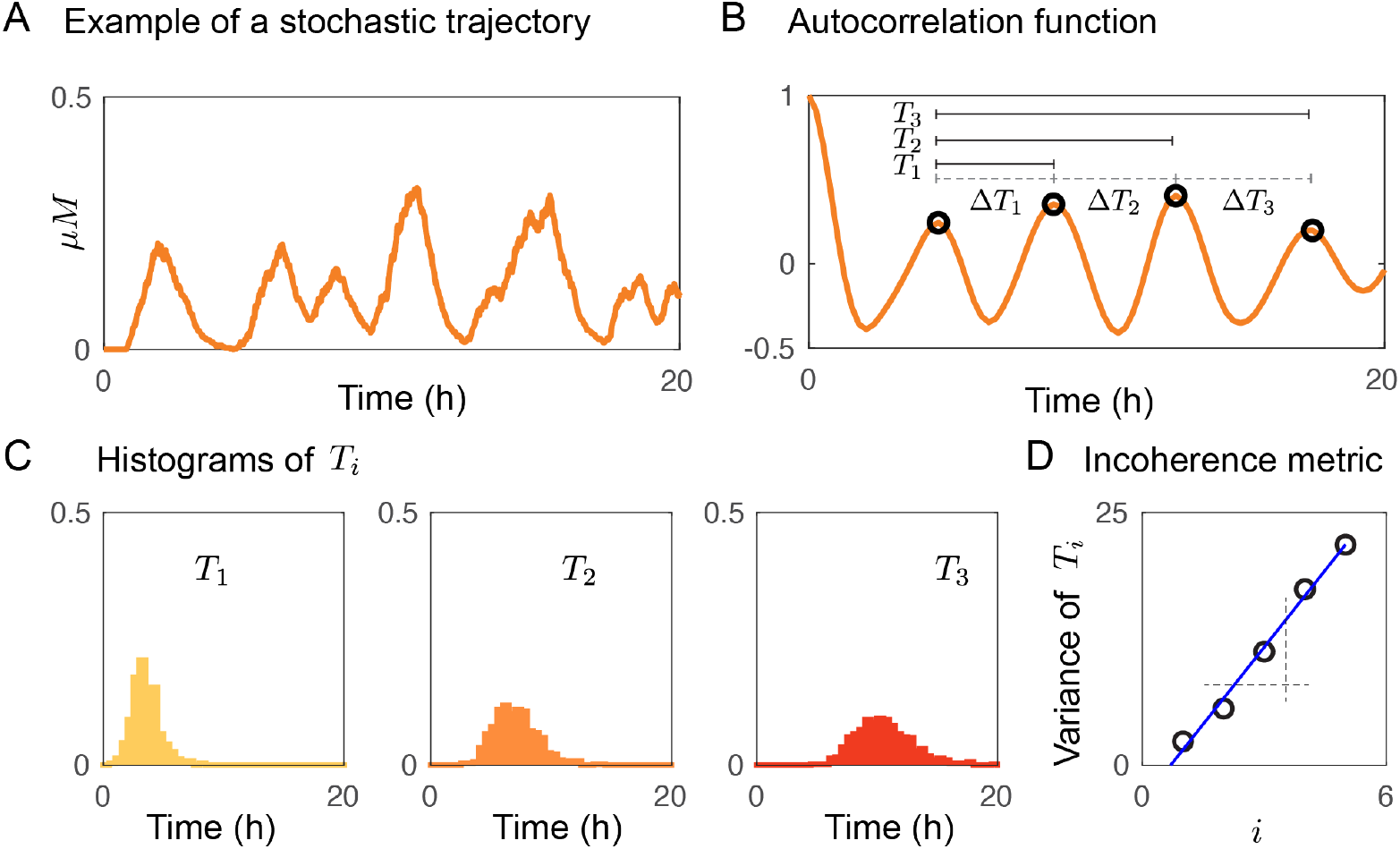
Incoherence metric for stochastic trajectories. **A)** Example of a stochastic trajectory with periodic behavior. **B)** The autocorrelation function of the orange plot in (A) highlights the periodic cycles from the stochastic trajectory. *T*_*i*_ denotes the time interval between peak 1 and peak i + 1, while the time interval between subsequent peaks *i* + 1 and *i* is denoted as Δ*T*_*i*_. **C)** Histograms showing the distributions of *T*_1_, *T*_2_, and *T*_3_, when their values are taken from multiple simulations. **D)** The variance of *T*_*i*_ over multiple simulations plotted against i can be approximated as a line, whose slope is a metric for incoherence.

Given a collection of stochastic simulations, the histograms of the inter-peak time intervals Δ*T*_*i*_ provide direct information on the period variability; coherence in period would be associated with similar mean and variance for all the Δ*T*_*i*_. It is more advantageous however to consider statistics of the *T*_*i*_ intervals, in particular their variance. While *T*_1_ = Δ*T*_1_, one expects the average of *T*_*i*_, *i* > 1, to increase linearly with *i*. While the variance of Δ*T*_*i*_, *i* > 1, should not increase if oscillations are coherent, the variance of *T_i_* does linearly increase, as fluctuations in each period are added (over each cycle) in the computation. The variance should increase proportionally to the level of “incoherence” of the oscillations. Thus, we focus on the variance of the histograms of the *T*_*i*_ intervals as exemplified in Fig. 1C. The variance of each *T*_*i*_ histogram is then plotted against the peak index, as shown in Fig. 1D, and the slope of this plot is used as an *incoherence metric* for the period. The variance tells us how regular the period is because a high variance means there is a large variety of periods. In other words, these values are irregular when variance is high. Low variance in period is required for coherent oscillations, which is the case when the metric is low. This metric was computed with our simulations of the six different designs in different parameter regimes to compare their robustness to changes in the various parameters, including recombinase translation rate constant and degradation rate constant.

## 3 Results

### 3.1 Building a biomolecular oscillator by coupling two self-inhibiting recombinases

Our basic design for achieving a periodic behavior using recombinases is shown in Fig. 2A. It consists of a single promoter between two recombinase binding sites, that controls the expression of two genes encoding distinct recombinase homodimers: *X*_1_, a serine integrase that targets attP and attB binding sites, and *X*_2_, the same serine integrase, but fused to its recombination directionality factor (RDF), which targets attR and attL binding sites (see Box 1). The expected operation of the circuit is the following: when the promoter points to the left, with the attP and attB binding sites on either side, it produces the recombinase *X*_1_; when the level of *X*_1_ is sufficiently high, it causes inversion of the promoter, pointing it to the right. In this way, *X*_1_ inhibits its own production. When the promoter points to the right, then it has attR and attL binding sites on either side and it allows for production of recombinase *X*_2_; in turn, *X*_2_ causes inversion and return of the promoter to the left-pointing orientation, thereby inhibiting its own production. The coupling of these two self-inhibiting modules is expected to periodically switch the promoter between the left-pointing and right-pointing configurations. This has been mathematically proved in a deterministic scenario [24]. Since a potential problem with this design is the accumulation of recombinase proteins, we also introduce a sequestration reaction between *Z*_1_ and *Z*_2_, the monomers forming *X*_1_ and *X*_2_, respectively. Serine recombinases can realize sequestration through heterodimerization, which promotes the removal of the non-limiting species [36].

**Figure 2:**
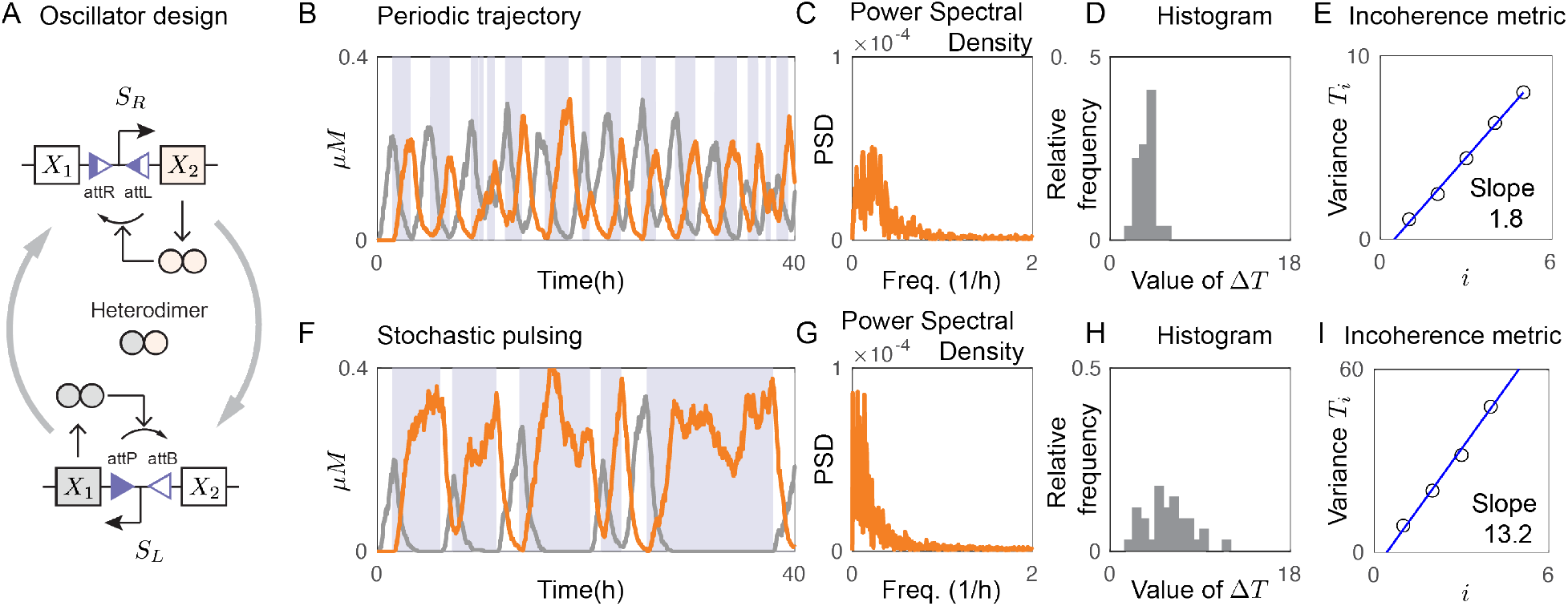
Coherence of a recombinase-based oscillator. **A)** Recombinase-based oscillator design consisting of two coupled, self-inhibiting modules. Recombinases X_1_ and X_2_, each inverting the promoter that controls its own production, create a tug-of-war-like behavior. The binding sites on either side of the promoter change back and forth at each inversion. **B)** Example of a periodic trajectory achieved using the parameters in Table 1: x_1_ is in gray and x_2_ in orange. Light gray regions mark when the promoter points to the right (configuration *S*_*R*_) and white regions mark when it points to the left (configuration *S*_*L*_). **C)** Power Spectral Density of the trajectory in panel B. **D)** Distribution of the autocorrelation inter-peak times Δ*T*_*i*_ for the trajectory in panel B. **E)** Incoherence metric: slope of the line interpolating the variance of times *T*_*i*_ as a function of the peak index *i*, computed over an ensemble of 500 simulations (cf. Fig. 1D). **F)** Example of a trajectory exhibiting stochastic pulsing, with corresponding **G)** Power Spectral Density, **H)** inter-peak period histogram, and **I)** incoherence metric plot.

To computationally characterize the behavior of this circuit, we considered a set of chemical reactions that model transcription, translation, and sequestration interactions between recombinase monomers, and promoter inversion. The rates of these reactions were converted to reaction propensities, expressing the probability of a reaction event per unit time, and we used the Gillespie algorithm to simulate the system (see Methods). Depending on its orientation, the promoter regulates the transcription of either mRNA *M*_1_ (when it points to the left in configuration *S*_*L*_) or mRNA *M*_2_ ( when it points to the right in configuration *S*_*R*_), with rate constant *θ*. Both mRNAs are assumed to dilute/degrade with a rate constant *ϕ*. In addition, the mRNAs *M*_1_ and *M*_2_ respectively are translated to recombinase monomers *Z*_1_ and *Z*_2_ with rate constant *ρ*.

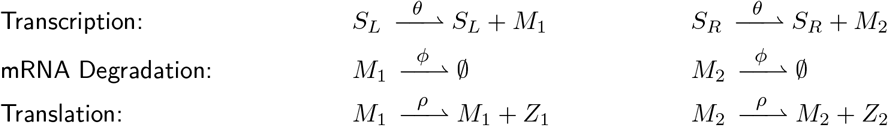

The serine recombinase monomers *Z*_1_ and *Z*_2_ can form the homodimers *X*_1_ and *X*_2_, respectively. Since *Z*_2_ is *Z*_1_ fused to its RDF, the two species can also form the heterodimer *C*. For simplicity we assume all dimers have an association rate constant *a* and a dissociation rate constant *d*. In addition, we assume all proteins degrade with a rate constant *δ*.

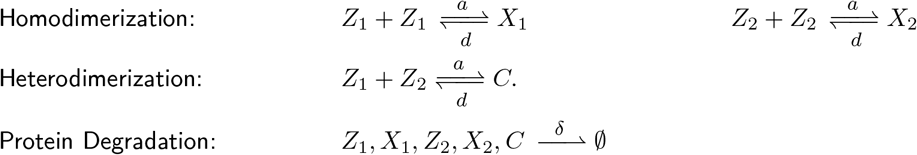

The rate of promoter inversion is regulated by the recombinase dimers *X*_1_ and *X*_2_ with rate parameters *f*_1_ and *f*_2_.

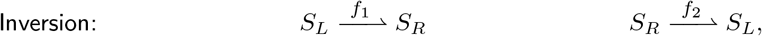

where 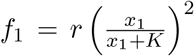 and 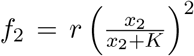, with *r* and *K* positive constants. The derivation of expressions *f*_1_ and *f*_2_ can be found in the STAR methods.

### 3.2 Coherence of stochastic simulations

We explored the emergence of periodic behaviors in the design described above via stochastic simulations, and in particular we wished to assess whether the circuit supports the occurrence of oscillatory solutions with a regular period. Fig. 2B shows example trajectories of recombinase concentration computed using the parameters from Table 1: *X*_1_ is in gray and *X*_2_ in orange. The light gray regions mark when the promoter points to the right (configuration *S*_*R*_). The trajectories for *X*_1_ and *X*_2_ show oscillations with anti-phase behavior (one level increases while the other decreases). We plotted the Power Spectral Density (PSD) of the orange trajectory (Fig. 2C) and detected the presence of a narrow interval of dominant frequencies, which is a hallmark of periodic behavior. Yet, since stochastic noise makes it challenging to identify a defined period, we examined the coherence of periodicity by computing the trajectory’s autocorrelation and then calculating the inter-peak times Δ*T*_*i*_ of the autocorrelation function (cf. Fig. 1B). We generated a histogram with the inter-peak times Δ*T*_*i*_ (Fig. 2D) that shows low variability of the period. As a metric to evaluate the incoherence of oscillations, we measured the time from the first peak of the autocorrelation function to six consecutive peaks in 500 trajectories. A small variance for these time intervals corresponds to trajectories with a small amount of change, while higher variance indicates oscillations with more change in period over time, but anyway the variance increases approximately linearly with the peak index (cf. Fig. 1D). The variance of the distribution of the times *T*_*i*_ was plotted against the peak index *i* (Fig. 2E): we use the slope of this line as a metric to quantify the incoherence of oscillations (see Methods). The smaller is the slope, the more coherent are the oscillations.

While the circuit design in Fig. 2A can exhibit coherent oscillations (as shown in Fig. 2B for the parameters in Table 1), it can also yield switching behaviors with significant period variability when parameters deviate from the nominal values reported in Table 1. By lowering the maximum switching rate to *r*/5, for example, the system’s switching behavior becomes less consistent. This means that promoter inversion becomes a random event, and we observe a regime that we call stochastic pulsing, illustrated with an example simulation in Fig. 2F. Stochastic pulsing is characterized by high variance in period (and possibly low variance in amplitude). The corresponding Power Spectral Density (Fig. 2G) lacks a clear dominant frequency, and the histogram of the inter-peak times of the autocorrelation function shows a period distribution with higher variance (Fig. 2H). Also, Fig. 2I shows how fast the variance of *T*_*i*_ increases: the slope is 13.2 (while it is 1.8 for the example of periodic trajectory), thus confirming the effectiveness of the used incoherence metric. The irregular frequency of switches in the promoter position can be noticed in Fig. 2F by looking at the irregularly-spaced, light gray bands, marking when the promoter is pointing to the right.

### 3.3 Analysing a single self-inhibiting mechanism to understand the coherence of oscillations

We sought to explain more clearly the possible emergence of incoherent oscillations in the circuit design in Fig. 2A. Because the design consists of two interconnected self-inhibitory modules, we reasoned that a single self-inhibitory module contributes to half of each oscillatory cycle. We examined the behavior of a self-inhibitory module by computing the time until inversion occurs, denoted as *T*_*I*_, and the recombinase concentration at inversion, denoted as *X*_*I*_. Both quantities have statistical properties that make it possible to elucidate the impact of noise propagation on the switching behavior. Further, the analysis of *T*_*I*_ and *X*_*I*_ contributes to the identification of key network parameters that mitigate the phase change in the oscillations.

We focus on the chemical reactions modeling a single self-inhibiting module (see the schematic in Fig. 3A):

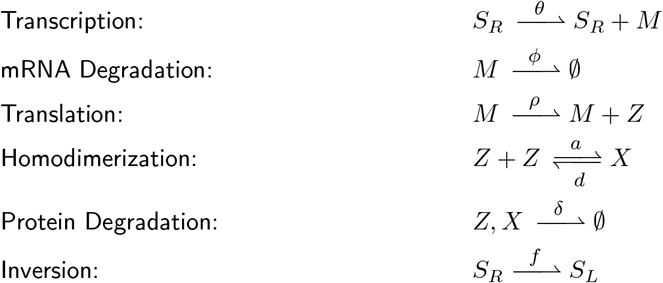

where 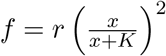, *K* > 0 and parameter *r* > 0 is the maximal switching rate.

**Figure 3:**
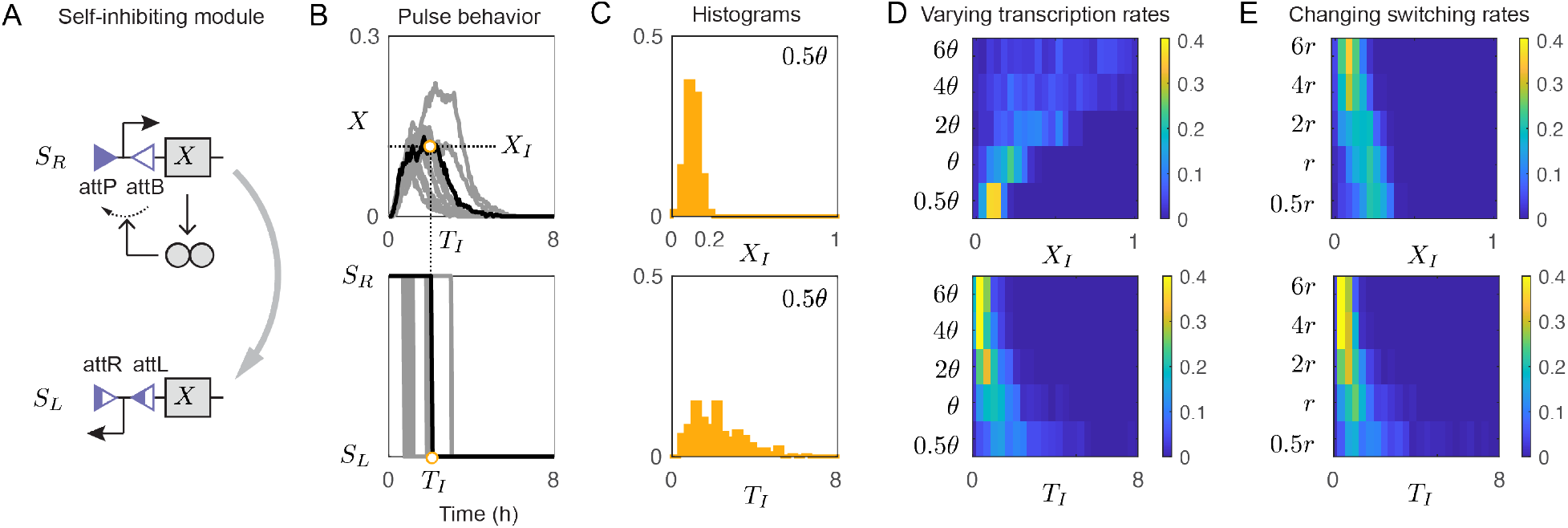
Analysis of the self-inhibiting module. **A)** Schematic describing the operation of the self-inhibiting module. Top: The promoter points to the right (configuration *S*_*R*_) and thus allows for the production of recombinase **X**, which inverts the promoter. Bottom: After inversion, the promoter points to the left (configuration *S*_*L*_) and the recombinase **X** is no longer produced. **B)** Top: 10 example stochastic trajectories of the self-inhibiting module. All the trajectories show a pulse-like behavior, because the concentration of **X** increases until the promoter is inverted and the production of **X** stops, and at that point the level of **X** decays due to dilution/degradation. Bottom: Promoter position over time for each simulation. For all the simulations, the promoter initially points to the right (configuration *S*_*R*_) and eventually points to the left (configuration *S*_*L*_). The inversion time is denoted as *T*_*I*_ and the recombinase concentration at inversion is **X**_*I*_. **C)** For 500 simulations with a recombinase transcription rate of *θ*/2, the histograms show the relative frequency of two important quantities. Top: Histogram of recombinase concentration at inversion, *X*_*I*_; this variable has a low variance. Bottom: Histogram of the inversion time, *T*_*I*_; here we observe a high variance. **D)** Top: Heat map where each row represents a histogram of the recombinase concentration at inversion, *X*_*I*_, for 500 simulations with a different transcription rate **θ**. Bottom: Heat map where each row represents a histogram of the inversion time, *T*_*I*_, for 500 simulations with different transcription rates *θ*. **E)** Top: Heat map where each row represents a histogram of the recombinase concentration at inversion, *X*_*I*_, for 500 simulations in which the maximum switching rate *r* is varied. Bottom: Heat map where each row represents a histogram of the inversion time, *T*_*I*_, for 500 simulations, each with a different *r* value.

The self-inhibitory module produces a pulse in recombinase concentration (half of an oscillatory cycle) when the promoter is in the on-state (*S*_*R*_), until the level of recombinase is sufficiently high to elicit inversion of the promoter to the off-state (*S*_*L*_). When the promoter is off, the recombinase level decreases due to dilution/degradation. All stochastic simulations produce a pulse, as shown by the gray trajectories in Fig. 3B (top). Each trajectory presents a distinct promoter inversion time *T*_*I*_ and recombinase concentration *X*_*I*_ at the point of inversion; we mark these quantities in the figure for the trajectory in black. The top and bottom panels of Fig. 3C respectively show the histograms of the relative frequency of *X*_*I*_ values and of *T*_*I*_ values for 500 simulations, with a transcription rate constant *θ*/2. While *X*_*I*_ has small variance, *T*_*I*_ has a very large variance, indicating that the half-cycle of an oscillator using this particular transcription rate would have an irregular period. Thus, many consecutive, irregular half-cycles could cause the oscillator to get out of phase very quickly.

Next, we sought to gain a more comprehensive understanding of how the histograms of *X*_*I*_ and *T*_*I*_ are affected by the network parameters. Fig. 3D and Fig. 3E show heat maps representing the computed distribution of *X*_*I*_ (top) and *T*_*I*_ (bottom); each row of these plots corresponds to a different transcription rate *θ* (panel D) or switching rate parameter *r* (panel E). Higher transcription rates reduce both the mean and the variance of *T*_*I*_, but introduce a high variance for *X*_*I*_. When two such self-inhibiting modules are interconnected in the oscillator design in Fig. 2A, the resulting system exhibits rapid switching, which may not be coherent. In contrast, small values of *θ* lead to reduced variance for *X*_*I*_, but increased variance for *T*_*I*_. In the complete oscillator in Fig. 2A, such a low transcription rate would cause stochastic pulsing, because the inversion times have large variability. This observation based on Fig. 3D suggests that there is a window of recombinase production rates that yields regular oscillations. Still, even though coherence would improved with the right choice of the parameter values, because the variance would be lower in both the distribution for *X*_*I*_ and for *T*_*I*_, the suitable range of parameters would be very small, hence the oscillatory behavior would be fragile and would not exhibit robustness even with respect to very small parameter variations. A similar effect can be observed if we change the translation rates and the degradation rates of mRNA and proteins, because they also affect the switching rate. In Fig. 3E, we varied the maximal switching rate parameter *r*, which scales the inversion propensity *f* of the promoter. Overall, a small parameter *r* leads to high variance in the distribution of *T*_*I*_, and hence the trajectories of the corresponding full oscillator design in Fig. 2A exhibit stochastic pulsing. In contrast, a large switching parameter *r* results in small mean and variance for both *T*_*I*_ and *X*_*I*_, thus improving the coherence of oscillations. However, extremely large values of *r* may induce very rapid switching in the complete circuit.

Overall, this analysis indicates that a characterization of the self-inhibiting module in isolation can provide insights on the operation of the full oscillator, when two self-inhibiting modules are suitably interconnected as in Fig. 2A. Our computations elucidate the effect of two parameter values on the mean and variance of relevant features of the pulsing behavior (for the individual self-inhibiting module), and connect those features to the likelihood of yielding coherent oscillations in the full circuit design (when two self-inhibiting modules are interconnected). In the next sections, we assess whether structural changes in the reaction network may expand the parameter range for which coherent oscillations occur.

### 3.4 Architectures improving the coherence of oscillations

In the previous section, we proposed and examined a recombinase circuit design that can exhibit a periodic switching behavior. In this section, we explore systematically how we can improve the coherence of the oscillations by modifying specific parameters as well as by modifying the circuit architecture. We focus in particular on the effects of adding different reaction mechanisms: repression, and catalytic degradation.

First, we note that sequestration of recombinases is present in all our designs, as we are using serine integrases. After an inversion event, mutual sequestration between the recombinases depletes both recombinase species, so that the next inversion cannot occur until the level of recombinase being expressed exceeds the level of recombinase produced at the previous cycle. This means that more time is needed before the next switching event occurs, relative to when sequestration is absent. Because preliminary analysis of the circuit in the absence of sequestration did not yield coherent oscillations, we reasoned that the circuit benefits from mechanisms that should space out in time the switching events. We begin by considering a design that includes repressors of recombinase transcription (RR design) to control more tightly the level of recombinase after inversion, reducing the potential for low copy fluctuations of the recombinase level. Similarly, the removal of recombinases via catalytic degradation would also control more tightly noise at low levels of recombinase expression. Thus, we consider a circuit design including the proteolytic degradation of recombinases (RP design). Both transcriptional repression and protease degradation, by reducing the available recombinase, have the result of delaying the next switching event [37]. We further explore two additional designs: one that incorporates sequestration of recombinase mRNA via complementary small-RNA like (sRNA) species that prevent translation (RS design), and another that regulates recombinase concentration by means of a transcriptional activator (RA design). Both these additional reactions can lengthen the switching period, which has the potential to increase coherence of the oscillatory behavior. We used the Gillespie algorithm to generate an ensemble of trajectories for each circuit, starting with the recombinase-based oscillator we have already proposed (see Fig. 2A). We then assess the corresponding period coherence. In each case, we vary the nominal parameters within a given range.

#### 3.4.1 R design: analysing the coherence of the recombinase-based oscillator

The R design was preliminarly illustrated in Fig. 2A. Its schematic is provided in Fig. 4A, along with sample trajectories, so as to ease comparison with possible alternative circuit designs. In Fig. 4D, we comparatively analyze the circuit behavior and we examine the coherence of the resulting oscillations as multiple parameters are varied. A lower translation rate constant, *ρ*, results in lower recombinase concentration, which leads to stochastic pulsing as reflected in the higher values of the incoherence metric in the orange plot in Fig. 4D (1). Analogously, a lower transcription rate *θ* gives rise to a higher variance in *T*_*I*_, as we have seen in Fig. 3D, bottom. The orange plot in Fig. 4D (1) also shows that a higher translation rate reduces the incoherence metric for this circuit, thus indicating improved coherence of oscillations. However, it is worth keeping in mind that very high values of *ρ* can lead to very rapid switching.

**Figure 4:**
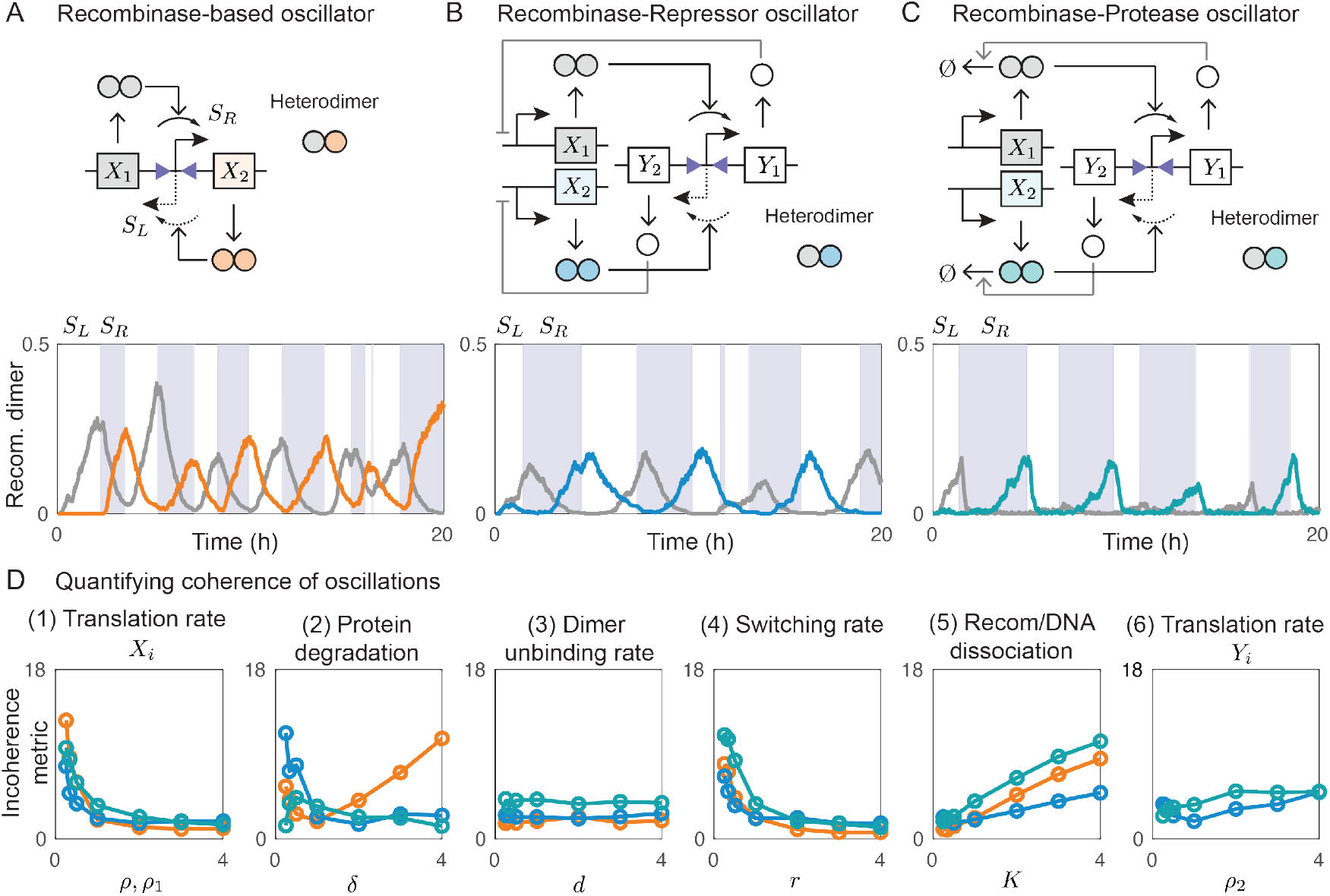
Analysis of the R, RR, and RP oscillator designs. **A)** Top: R oscillator design with a single inverting promoter that alternately controls the production of recombinases *X*_1_ (when it is pointing to the left, configuration *S*_*L*_) and *X*_2_ (when it is pointing to the right, configuration *S*_*R*_). Molecular sequestration is included through the heterodimerization of *X*_1_ and *X*_2_. Bottom: Trajectories of a single simulation showing the time evolution of the concentrations of *X*_1_ (gray) and *X*_2_ (orange) over time. The light gray stripes mark when the promoter points to the right (configuration *S*_*R*_), while white stripes mark when the promoter points to the left (configuration *S*_*L*_). **B)** Top: RR oscillator design with a single inverting promoter that alternately controls the production of two different repressor proteins, *Y*_1_ and *Y*_2_, while recombinases *X*_1_ and *X*_2_ are produced constitutively and also heterodimerize. Bottom: Trajectories of a single simulation showing the time evolution of the concentrations *X*_1_ (gray) and *X*_2_ (blue). The colored stripes indicate the current promoter configuration (*S*_*R*_ or *S*_*L*_). **C)** Top: RP oscillator design with an inverting promoter that alternates between producing protease proteins *Y*_1_ and *Y*_2_, while recombinases *X*_1_ and *X*_2_ are produced constitutively with the ability to heterodimerize. Bottom: Trajectories of a single simulation showing the time evolution of *X*_1_ (gray) and *X*_2_ (teal). The colored stripes denote the current promoter configuration (*S*_*R*_ or *S*_*L*_). **D)** Analysis of the coherence of the R, RR, and RP designs for different parameter regimes. Each point on these plots represents the incoherence metric calculated using a collection of simulations using the parameter value indicated on the x-axis. All other parameters used for our simulations are reported in Table 1 along with their nominal values. For our sensitivity analysis, in each plot we vary the considered parameter value from 0.25 to 4 times its nominal value.

Increasing the degradation rate, *δ*, can also causes stochastic pulsing by reducing the amount of recombinase in the system to very low levels, as is reflected in the high value of the incoherence metric in the orange plot in Fig. 4D (2). Low degradation *δ* also corresponds to a high value of incoherence metric in the orange plot of Fig. 4D (2), since it could lead to a high recombinase concentration and therefore to rapid switching. Overall, our incoherence metric indicates there is a range of *δ* values that yield strong coherence of oscillations, while very low or very large values hinder coherence.

The incoherence metric remains fairly constant in the wide range of *d* values shown in the orange plot of Fig. 4D (3), which means that coherence is unaffected by reasonably small changes in the value of the dimer dissociation rate. However, decreasing *d* by a factor of 100 does affect the incoherence metric (not shown).

A low switching rate, *r*, is associated with a high incoherence metric, and we observed this results in stochastic pulsing because the probability of inversion is lower. Conversely, increasing the value of *r* improves the coherence of this design (the incoherence metric is much lower), as can be seen in the orange plot in Fig. 4D (4). However, higher values of *r* may lead to very fast switching, which may be too rapid to be classifiable as a suitable oscillation.

Higher values of *K* mean that switching requires larger amounts of recombinase. Thus, a large *K* increases irregularity of switching, and therefore reduces coherence, because the requirements for inversion are not met and yet the promoter is induced to invert anyway. On the other hand, with smaller values of *K*, a lower amount of recombinase is required for the inversion to occur, and hence the system can approach the maximum switching rate due to saturation, leading to an overall lower incoherence metric, as shown in the orange plot in Fig. 4D (5).

#### 3.4.2 RR design: transcriptional repression of recombinase production

The RR design, whose schematic is shown in Fig. 4B along with an example trajectory, incorporates transcriptional repressors as a form of cascaded regulation. In this design, the transcription of recombinases *X*_1_ and *X*_2_ is regulated by transcriptional repressors *Y*_1_ and *Y*_2_, respectively. The production of *Y*_1_ and *Y*_2_ is controlled by an inverting promoter positioned between the recombinase binding sites (as for the production of recombinases in the R design), while the genes for *X*_1_ and *X*_2_ have promoters regulated by *Y*_1_ and *Y*_2_, respectively, acting as repressors. When the promoter of the genes points to the right, *Y*_1_ is produced, inhibiting the production of recombinase *X*_1_. At the same time, the amount of recombinase *X*_2_ increases because *Y*_2_ is not being produced. This causes the promoter to invert to the left. As a consequence, the production of *Y*_2_ increases, while *Y*_1_ decreases in concentration due to decay. The remaining *Y*_1_ still blocks the production of *X*_1_ until its level is too low for repression. When *Y*_2_ is high in concentration and *Y*_1_ is low, the amount of recombinase *X*_1_ increases and eventually becomes sufficient to invert the promoter to the right. In addition, *Y*_2_ delays the production of *X*_2_ until the concentration of *Y*_2_ becomes low enough, making it unable to repress the production of *X*_2_. This sequence of steps is expected to repeat as the promoters invert back and forth. Overall, the repressors introduce a lag in the response time of the system, because each repressor protein remains present after promoter inversion and continues to repress its associated promoter effectively. Additional time is required for the repressor to decay to levels where it no longer represses the production of its associated recombinase. The sequestration reaction between recombinases further helps keep the less abundant recombinase species inactive, thereby preventing stochastic pulsing and promoting more coherent oscillations.

We developed a model for the RR circuit (Fig. 4B), reported here as a list of chemical reactions including transcription and translation. Rate constants were converted to propensities to simulate the reactions using the Gillespie algorithm. We then studied the periodic behavior of this circuit under different parameter values.

The promoter regulates the transcription of either of the two mRNA repressors, *W*_1_ and *W*_2_, with rate constant *θ* at any given time. *W*_1_ is produced when the promoter is pointing to the right (configuration *S*_*R*_) and *W*_2_ is produced when the promoter is pointing to the left (configuration *S*_*L*_). The mRNAs also decay with a rate constant *ϕ*.

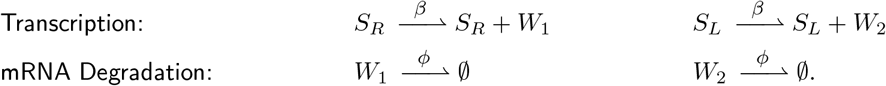

*W*_1_ and *W*_2_ are translated into repressors proteins *Y*_1_ and *Y*_2_ with a rate constant *ρ*_2_, and decay with a rate constant *δ*.

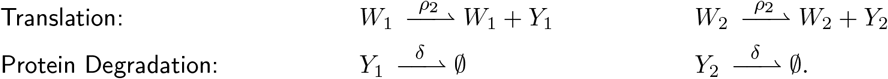

The repressors *Y*_1_ and *Y*_2_ regulate the transcription rate of recombinase mRNAs *M*_1_ and *M*_2_, which are produced with rate parameters *g*_1_ and *g*_2_, respectively, given by expressions for Hill-type repression of dimers. Both mRNAs decay with a rate constant *ϕ*. In addition, the mRNAs produce recombinase monomers *Z*_1_ and *Z*_2_ with a rate constant *ρ*_1_.

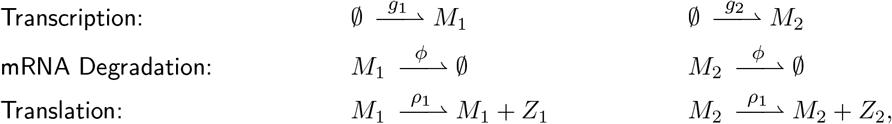

where 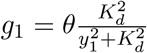 and 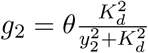.

The monomers *Z*_1_ and *Z*_2_ form homodimers *X*_1_ and *X*_2_ in addition to heterodimer *C*, both with an association rate constant *a* and a dissociation rate constant *d*. The heterodimer formation is possible because the second recombinase monomer is fused to its RDF. In addition, all proteins complexes decay with a rate constant *δ*.

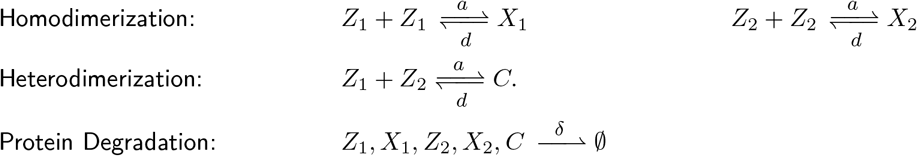

Finally, the switching rate of the promoter is regulated by the recombinase dimers *X*_1_ and *X*_2_ with rates *f*_1_ and *f*_2_:

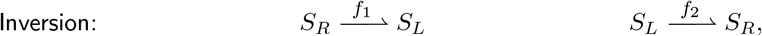

where 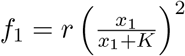 and 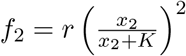.

As shown in Fig. 4D (1), blue plot, the RR design has a high incoherence metric for low values of translation rate constant, *ρ*_1_, while coherence increases for higher *ρ*_1_ values, as it happens for the R design. Similarly, high *ρ*_1_ can lead to fast switching and low *ρ*_1_ can give rise to stochastic pulsing. Fig. 4D (2), blue plot, shows that, compared to the R design, the RR design has a wider range of *δ* values for which the incoherence metric is low, therefore yielding regular oscillations. This is a noticeable improvement in performance between designs. As for the R design, coherence is unaffected by *d*, as shown in the blue plot in Fig. 4D (3), unless *d* is decreased by a factor of 100 (not shown). This design also has a high incoherence metric for low values of the switching rate, *r*, while the incoherence metric gets much smaller when *r* is high, as shown in Fig. 4D (4), blue plot. The blue plot in Fig. 4D (5) shows that, as for the R design, the incoherence metric becomes higher as the *K* value of the RR design increases, revealing poor coherence between periods, because promoter inversion is induced only at too high recombinase concentrations. There is also a small increase in the incoherence metric as the repressor translation rate, *ρ*_2_, increases, as shown in Fig. 4D (6), blue plot.

#### 3.4.3 RP design: introducing protease-based regulation

Since slowly-decaying recombinases lead to higher rates of inversion for the R design (cf. Fig. 4D (2), orange plot), we reasoned that regulating the decay rate could improve the switching behavior of an oscillator. Next, we analyze a circuit design that relies on proteases to control the amount of active recombinase present. The circuit schematic is shown in Fig. 4C, along with an example trajectory. Proteases are proteins that can cleave a polypeptide at a specific, targeted amino-acid sequence so as to inhibit its function. [38, 39]. The RP circuit design consists of recombinases *X*_1_ and *X*_2_ that are cleaved by two orthogonal proteases *Y*_1_ and *Y*_2_, respectively. When the inverting promoter points to the right, it produces the protease *Y*_1_, which selectively cleaves the recombinase *X*_1_. At the same time, the amount of recombinase *X*_2_ increases due to the lack of protease *Y*_2_. Then *X*_2_ inverts the promoter to the left, initiating the production of protease *Y*_2_ and leaving *Y*_1_ to decrease in concentration due to cleavage. However, the remaining *Y*_1_ continues to target and cleave *X*_1_. This delays the increase in the amount of *X*_1_. After this delay, *X*_1_ eventually inverts the promoter to the right and stops the production of *Y*_2_. The remaining amount of *Y*_2_ will delay the production of recombinase *X*_2_ before it can then effectively invert the promoter to the left. These interactions are expected to generate a repetitive switching behavior in which the promoter is inverted from right to left back and forth, but the half-cycle of each oscillation is expected to be longer when compared to the R design.

As in the previous cases, we built a model to evaluate the dynamics of the RP circuit design shown in Fig. 4C. Constitutive transcription of recombinase mRNAs *M*_1_ and *M*_2_ occurs at a constant rate *θ*, and decay at a constant rate *ϕ*. In addition, the mRNAs produce recombinase monomers *Z*_1_ and *Z*_2_ at a constant rate *ρ*_1_.

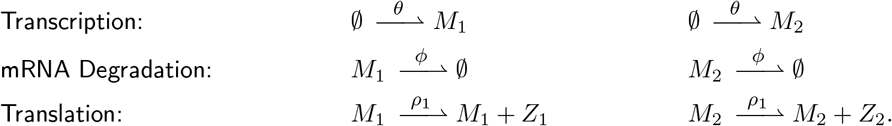

The monomers *Z*_1_ and *Z*_2_ can form homodimers *X*_1_ and *X*_2_ as well as heterodimer *C* with association rate constant *a* and dissociation rate constant *d*. The heterodimer formation is possible because the second recombinase monomer is fused to its RDF. In addition, all protein complexes decay at a constant rate *δ*.

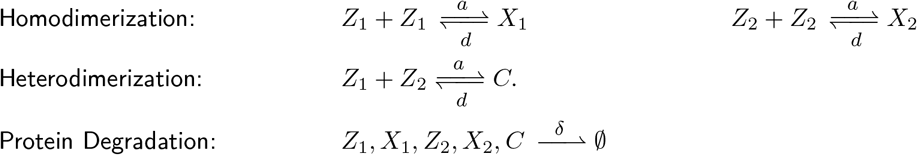

The inverting promoter, located between recombinase binding sites, regulates the transcription of protease mRNAs *W*_1_ and *W*_2_ at rate *β*. Protease mRNAs are also degraded/diluted at rate *ϕ*.

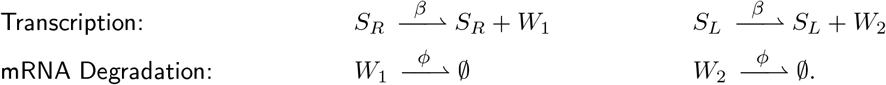

The mRNA species *W*_1_ and *W*_2_ yield proteases *Y*_1_ and *Y*_2_, each with a rate constant *ρ*_2_. The protease proteins decay with a rate constant *δ*.

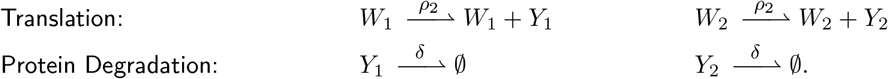

The protease *Y*_1_ (respectively *Y*_2_) targets the recombinase monomer *Z*_1_ (respectively *Z*_2_) as well as dimer *X*_1_ (respectively *X*_2_) for catalytic degradation. These reactions occur with rate *g*_1_ (respectively *g*_2_), corresponding to a Hill-type function with Hill coeffiecient equal to 1, as derived in the STAR methods.

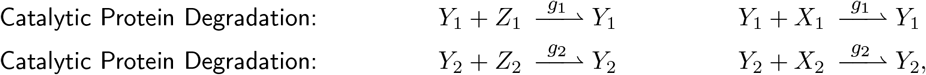

where 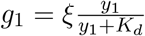 and 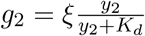.

Finally, the switching rate of the promoter is regulated by the recombinase dimers *X*_1_ and *X*_2_, with rates *f*_1_ and *f*_2_ as shown.

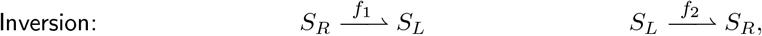

where 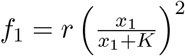 and 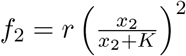.

Like the previous designs, the incoherence metric of the RP design is high at low *ρ*_1_ values, and it decreases as *ρ*_1_ increases, as shown in Fig. 4D (1), teal plot. Hence, a large *ρ*_1_ improves the coherence of oscillations. In contrast with other designs, this case has a remarkably low incoherence metric for low degradation rate *δ*, as show in Fig. 4D (1), teal plot: this indicates that stochastic pulsing does not occur at low degradation rates. For this reason, this design is the most robust to changes in *δ*, among all the designs we considered so far. Again, varying *d* does not affect coherence (Fig. 4D (3), teal plot), unless *d* is decreased by a factor of 100. Just like the R and RR oscillators, for the RP design a low *r* is associated with a high incoherence metric, which decreases as *r* increases (Fig. 4D (4), teal plot). When *K* varies, the metric follows the same pattern as for the R and RR oscillators: the incoherence metric is low for small *K* values and increases with *K* (Fig. 4D (5), teal plot). Fig. 4D (6), teal plot, shows the effect of increasing the protease translation rate, *ρ*_2_: the incoherence metric increases slightly as *ρ*_2_ increases, following the same trend observed when varying *ρ*_2_ (repressor protein translation rate) in the RR oscillator.

#### 3.4.4 Alternative architectures

Here, we briefly describe three additional architectures where recombinase levels are regulated with different approaches. The derivation of the model for each circuit design is in the STAR Methods section.

##### An activator-based circuit design (RA design)

The RA circuit design incorporates an activator to regulate recombinase production. The corresponding schematic is reported in Fig. 5A, along with an example trajectory. The design consists of the transcriptional activators *Y*_1_ and *Y*_2_ that drive the production of recombinases *X*_1_ and *X*_2_. When the inverting promoter points to the left, it produces the activator *Y*_1_, driving the production of *X*_1_. These two steps in the cascade must occur before *X*_1_ can be produced, and hence slow down the production of the recombinase. Once *X*_1_ increases in concentration, it can invert the promoter to the right. This leads to the production of *Y*_2_ and stops the production of *Y*_1_. Then, *Y*_2_ increases the production of *X*_2_. When *X*_2_ increases in concentration, it can invert the promoter to the left. Overall, this leads to a switching cycle of inverting the promoter from left to right repeatedly. One challenge with this design is that, even when an activator is not being produced, the remaining transcription factor can still increase the production of its associated recombinase. This makes it difficult for the the concentration of each recombinase to get very low, which causes the promoter to invert more irregularly.

**Figure 5:**
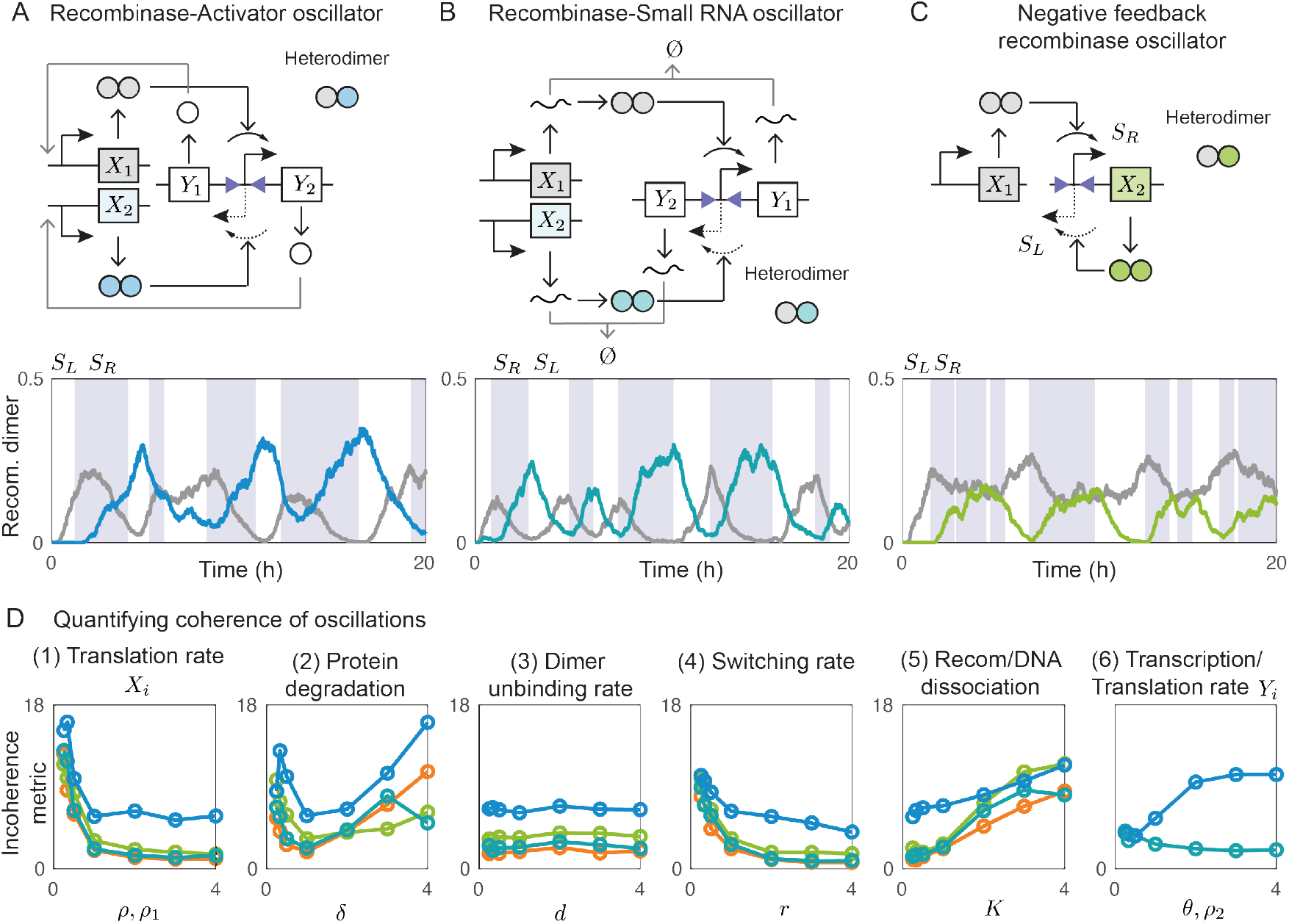
Analysis of the RA, RS, and NF oscillator designs. **A)** Top: RA oscillator design with a single inverting promoter that alternately controls the production of transcriptional activators *Y*_1_ and *Y*_2_, which respectively regulate the production of recomabinases dimers *X*_1_ and *X*_2_; the recombinase monomers undergo molecular sequestration through heterodimerization. Bottom: Trajectories of a single illustrative simulation showing the time evolution of the concentrations of *X*_1_ (gray) and *X*_2_ (blue) over time. The light gray stripes mark when the promoter points to the right (configuration *S*_*R*_) while white stripes mark when the promoter points to the left (configuration *S*_*L*_). **B)** Top: RS oscillator design with an inverting promoter that alternately regulates the production of small RNAs *Y*_1_ and *Y*_2_, which inhibit mRNA recombinases, preventing their transcription into *X*_1_ and *X*_2_, respectively. These recombinases are also able to heterodimerize. Bottom: Trajectories of a single simulation showing the time evolution of the concentrations of *X*_1_ (gray) and *X*_2_ (teal). The colored stripes indicate the current promoter configuration (*S*_*R*_ or *S*_*L*_). **C)** Top: NF oscillator design with a single self-inhibiting module controlling the production of *X*_2_ and a constitutive promoter controlling the production of *X*_1_, with recombinase monomers sequestering into heterodimers. Bottom: Trajectories of a single simulation showing the time evolution of *X*_1_ (gray) and *X*_2_ (green). The colored stripes denote the current promoter configuration (*S*_*R*_ or *S*_*L*_). **D)** Analysis of the coherence of the RA, RS, and NF designs for different parameter regimes. Each point on these plots represents the incoherence metric calculated using a collection of simulations, using the parameter value indicated on the x-axis. All other parameters used for our simulations are reported in Table 1 along with their nominal values. For our sensitivity analysis, in each plot we vary the considered parameter value from 0.25 to 4 times its nominal value.

As a result, the RA design is the worst-performing among the designs we tested. The incoherence metric is consistently higher than that of the other oscillators, in every parameter regime. When *ρ*_1_ is varied, the incoherence metric follows the same pattern as that of previous designs, decreasing as *ρ*_1_ increases (Fig 5D (1), blue plot). However, the incoherence metric is consistently higher than that of the other oscillators. The window of *δ* values for which this oscillator is more coherent (Fig. 5D (2), blue plot) may appear wider than that for the R oscillator (Fig. 4D (2), orange plot, also reported in Fig. 5 for the sake of comparison), but the incoherence metric never gets as low for the RA oscillator as it does for the R oscillator (albeit in a small region). Parameter *d* does not affect coherence in this design too, as shown by the blue plot in Fig. 5D (3), unless it is decreased by a factor of 100. While the incoherence metric does decrease as *r* increases, it again remains higher than for the other oscillators, over the entire range of *r* values we tested (Fig. 5D (4), blue plot). The incoherence metric increases for increasing *K* (Fig. 5D (5), blue plot), as for the other oscillator designs, but, differently from the case of the other oscillators, it is high also for low *K* values. Finally, the activator translation rate, *ρ*_2_, causes a significant increase in the incoherence metric as it increases (Fig. 5D (6), blue plot), a pattern noticeably different from that shown by all the other oscillators with an additional element in their cascade.

##### An sRNA-based circuit design (RS design)

Molecular sequestration can program temporal delays by setting concentration thresholds [40]. The time that the system takes to reach this threshold defines the delay caused by sequestration. We take inspiration from this mechanism to design a circuit that sequesters recombinase mRNA with small RNA as shown in Fig. 5B, which reports the schematic as well as an example trajectory. The circuit consists of small RNA molecules *Y*_1_ and *Y*_2_ that target the recombinase mRNAs *X*_1_ and *X*_2_ respectively. When the inverting promoter points to the right, it produces the small RNA *Y*_1_, which sequesters the recombinase mRNA *X*_1_. At the same time, the recombinase *X*_2_ increases in concentration, because no small RNA *Y*_2_ is present. This results in the inversion of the promoter to the left. Then, the amount of small RNA *Y*_2_ increases, while *Y*_1_ decays. Then, *X*_1_ increases and inverts the promoter to the right. This results in the continuous inversion of the promoter from left to right.

Overall, the RS design performs better than the RA design, but it is not as successful as the RR and or RP designs. The incoherence metric for varying *ρ*_1_ values (Fig. 5D (1), teal plot) is akin to that for the R, RR, and RP designs. Compared to the RA oscillator, the RS oscillator has a broader range of *δ* values for which the incoherence metric is low (Fig. 5D (2), teal plot). Like in all other designs, *d* does not affect coherence (Fig. 5D (3), teal plot), unless it is decreased by a factor of 100. The effect of *r* on this oscillator is also in line with the behavior for the R, RR, and RP designs: the incoherence metric decreases as *r* increases (Fig. 5D (4), teal plot). The incoherence metric is somewhat higher for the *K* values in the mid-range (Fig. 5D (5), teal plot), when compared to the R, RR, and RP designs, but it is still less impacted by this parameter than it happens for the RA design. Interestingly, the incoherence metric slightly decreases as *ρ*_2_, the transcription rate of the sRNA, increases (Fig. 5D (6), teal plot), suggesting that the presence of more sRNA improves the coherence of oscillations.

##### Negative feedback oscillator (NF design)

The NF design, whose schematic is shown in Fig. 5C along with an example trajectory, consists of a constitutive promoter controlling the production of recombinase *X*_1_ and a self-inhibitory module regulating recombinase *X*_2_. The presence of two distinct promoters differentiates this design from the circuit design in Fig. 4A, which includes a single promoter. The regulation (or orientation) of the promoter controlling the production of *X*_2_ experiences the suppression effect of negative feedback, strengthened by the heretodimer complex formation from recombinase monomers. This circuit can exhibit switching behavior with a consistent period, as well as stochastic pulsing behavior. When the total production rate of *X*_2_ is larger than that of *X*_1_, *X*_2_ not only inhibits its own production, by causing its promoter to point to the left, but it also limits the homodimer formation for *X*_1_. When *X*_2_ decays, the concentration of *X*_1_ increases, which inverts the promoter to the right configuration, *S_R_*. This results in periodic behavior with a consistent period. Conversely, stochastic pulsing occurs when the level of *X*_1_ is very low and the inversion of the promoter to the right occurs randomly. On the other hand, when the concentration of *X*_1_ is larger than that of *X*_2_, the system yields fast oscillations, because *X*_1_ is readily available to flip the promoter to the state *S_R_*. We quantify its switching behavior using the incoherence metric in Fig. 5D, shown in the green plots. The effect of the various parameters is akin to that observed for the other circuits, except for the fact that smaller values of the incoherence metric can be observed when the protein degradation, *δ*, is increased (Fig. 5D (2), green plot).

#### 3.4.5 Summary of simulation results

To quantify the effect of the parameters on the coherence of oscillations, we used our incoherence metric to assess how fast the variance of *T*_*I*_ increases with the peak index *i*. We found that the dilution/degradation rate parameter *δ* has a strong influence on the viability of our candidate oscillators; in particular, in all designs a small *δ* (slow degradation) is detrimental to achieving coherent oscillations, while a large *δ* (fast degradation) may have either beneficial or detrimental effects depending on the circuit architecture. The circuits including mechanisms to tighten the control of recombinase levels are insensitive to increases in *δ*. The R design is the simplest and is shown to achieve coherent oscillations, but it is particularly sensitive to *δ* and the coherence of oscillations degrades as this parameter increases; furthermore, this circuit requires large values of *ρ* and *r* and small values of *K* to yield sufficiently coherent oscillations. The RR and RP designs have expanded ranges of *δ* values yielding coherent oscillations, presumably due to the fact that the additional reactions slow down the response timescale of the system, but show similar patterns in the other parameters. All designs are essentially unaffected by changes in the dissociation rate of the dimers, *d*, unless the value is decreased by a factor of 100 or more. Overall, the RR, RP, and RS designs, which are the ones with an inhibitory element explicitly present in their cascade, are the most successful.

## 4 Discussion

We described and computationally modeled different molecular circuit architectures to induce periodic behaviors using recombinases. At the core of all our designs, two self-inhibiting loops are coupled through a target DNA site that undergoes periodic inversion (switching) cycles induced by recombinase expression. We examined six variants of this basic architecture that use various mechanisms to introduce tighter control of recombinase levels in the circuit with the goal of achieving regular oscillations. Each of these oscillators was examined using stochastic simulations, assuming that a single copy of each genetic component is present, capturing a realistic scenario for genomically integrated circuits. To evaluate the incoherence of oscillations, we used an approach that examines the statistics of the autocorrelation function of stochastic trajectories, and we used this metric to identify which kinetic rates improve oscillation coherence. Our simulations indicate that the designs combining recombinases and mechanisms to suppress or delay recombinase expression, such as sequestration, transcription repressors, and proteases, allow for an improvement in the coherence of oscillations. Overall, our results indicate that three key parameters make it possible to improve the coherence of oscillations: the threshold for recombinase operation, the recombinase production rate, and its degradation.

Throughout our computational analysis, we observed that some parameter combinations result in very rapid cycles of promoter inversion in which the peak levels of recombinase remain low yet irregular. This fast switching occurs in particular when the maximal switching rate and the dilution/degradation rates are large. When examining the autocorrelation function of trajectories that exhibit this fast switching, the distribution of inter-peak times shows a small variance, so the slope of the coherence plot remains low. This points to the fact that our coherence analysis cannot discriminate between a fast switching regime and slower, coherent oscillations.

When designing synthetic molecular oscillators, there is general consensus on the necessity of negative feedback, which can be destabilized via a high gain, delays, or positive feedback [9, 10, 11, 12, 13, 14]. Our circuit designs agree with this strategy, although our architecture is unique in that it couples two stable negative feedback loops through mutual activation, enabled by recombinase inversion of a target DNA site. Our computational simulations indicate that the parameters related to each individual self-inhibiting module influence the coherence of oscillations for the full circuit by determining the statistics of the peak amplitude and duration of each half-cycle (Figure3). We speculate that in experiments, these recombinase-based self-inhibiting modules may first be tuned individually, facilitating the synthesis and characterization of the complete oscillators. We also conjecture that nested clocks with multiple synchronized periods may be built by coupling different self-inhibiting components. While our computations cannot prove any structural robustness of the circuits we proposed, they suggest that periodic behaviors are likely to occur in a broad range of parameter values, supporting experimental realization of recombinase oscillators.

## 5 STAR Methods

### Resource availability

Stochastic simulations were implemented using custom scripts in MATLAB using a workstation with the following specifications: 8 GB RAM and a 2.9 GHz, dual-core Intel Core i5. Scripts are available upon request from the Lead Contact Elisa Franco

### Materials availability

This study did not generate new materials or reagents.

## Supplementary results and figures

### Derivation of the recombinase inversion model

In this section, we derive a mechanistic model describing the process of DNA inversion mediated by a recombinase. We will focus on the inversion process (Fig. 6A); a similar approach can be used for the reverse-inversion process. Fig. 6B shows the multiple steps we expect to occur toward inversion, and we include all possible interactions between recombinase dimers and the target binding sites. Our goal is to reduce these steps to a single equivalent inversion reaction, whose rate can be converted to a propensity for the reaction to occur in our stochastic simulations.

**Figure 6:**
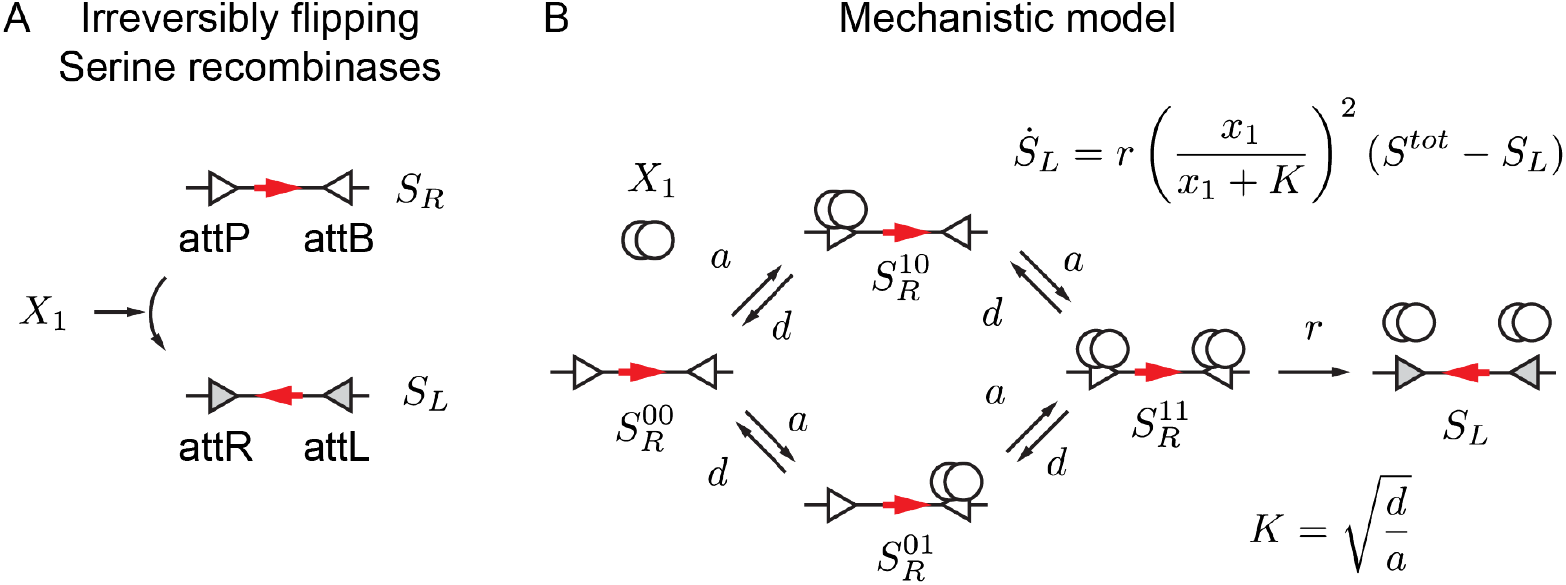
A: Schematic of the overall serine recombinase inversion process. B: Reactions steps required for inversion.

We write down an equivalent set of chemical reactions representing the steps shown in Fig. 6B; for simplicity, all association and dissociation constants are assumed to be the same.

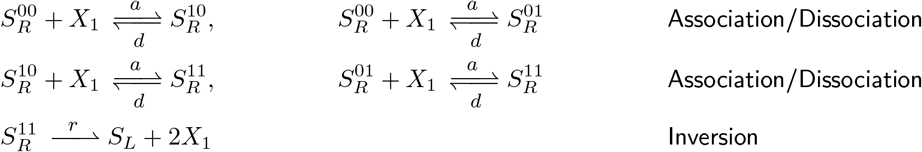

where 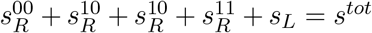; we denote the concentration of a chemical species *A* with the corresponding lowercase letter *a*. Then, we can write the ODEs as:

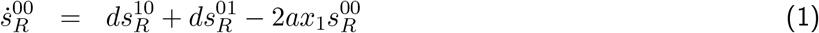

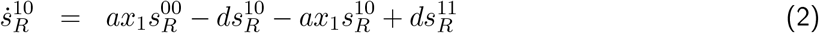

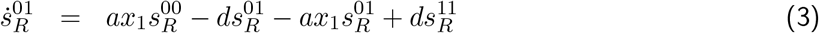

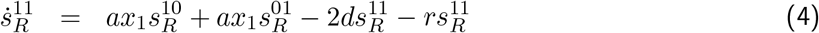

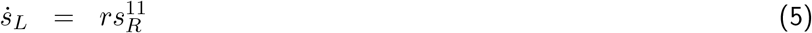

We assume that the association and dissociation constants are sufficiently fast so that the dynamics of 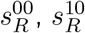 and 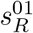 reach steady state rapidly, i.e. 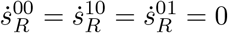. From 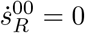, we find that

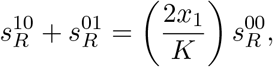

with *K* = *d/a*. From 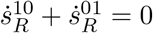, we get

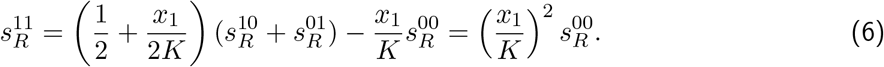

Then, we use the fact that the total amount of DNA is conserved:

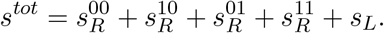

In the above equality, we can substitute the equilibrium level of 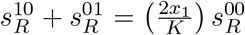, and then derive 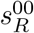 which can be substituted in Equation (6), finding:

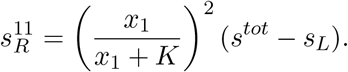

Finally, Equation (5) can be rewritten as follows:

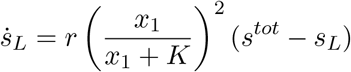

Next, we lump the intermediate states 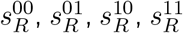 into a single variable 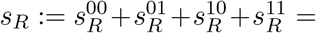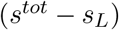. The dynamics of this variable are:

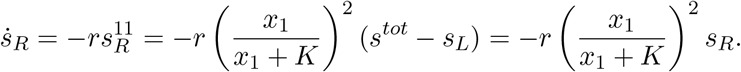

Therefore, in our stochastic simulations we simulate the promoter configuration *s*_*R*_ as a species that is inverted with propensity 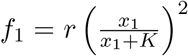 to the promoter in configuration *s*_*L*_. A similar reasoning can be carried out for the inversion from *s*_*L*_ back to *s*_*R*_, with a similarly defined propensity 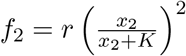, where again for simplicity we consider an identical dissociation constant and maximal conversion rate.

### RA design: introducing an activator-based cascade

We add a cascade that regulates the production of recombinases with transcriptional activators. The circuit schematic, shown in Fig. 5A, has an inverting promoter between recombinase binding sites that regulates the transcription of activator mRNAs *W*_1_ and *W*_2_ at a constant rate *β*. The mRNAs decay at a constant rate *ϕ*.

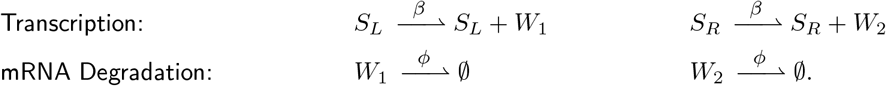

These mRNAs *W*_1_ and *W*_2_ produce repressor proteins *Y*_1_ and *Y*_2_ with a rate constant *ρ*_2_. These proteins decay with a rate constant *δ*.

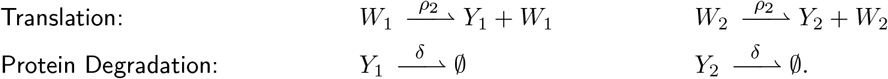

The activators *Y*_1_ and *Y*_2_ regulate the transcription of *M*_1_ and *M*_2_ at rates *g*_1_ and *g*_2_ (Hill-type repressor expressions for dimers), and both mRNAs decay with a rate constant *ϕ*. In addition, the mRNAs produce recombinase monomers *Z*_1_ and *Z*_2_ with a rate constant *ρ*_1_.

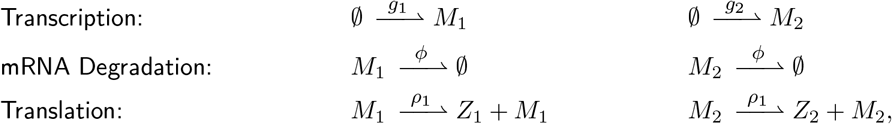

where 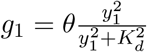 and 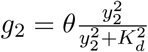.

The monomers *Z*_1_ and *Z*_2_ can form homodimers *X*_1_ and *X*_2_ and the heterodimer, *C*, with an association rate *a* and a dissociation rate *d*. The heterodimer formation is possible because the second recombinase monomer is fused to the RDF. In addition, all protein complexes decay with a rate constant *δ*.

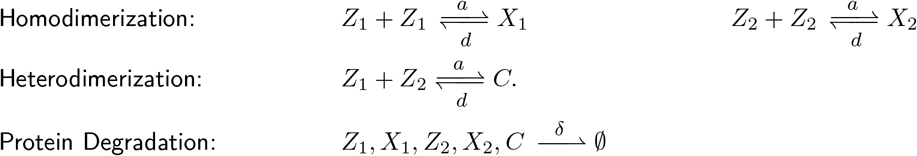

Finally, the inversion rate of the promoter is regulated by the recombinase dimers *X*_1_ and *X*_2_ with rates *f*_1_ and *f*_2_.

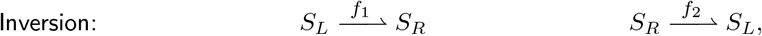

where 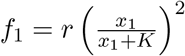 and 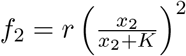.

### RS design: introducing recombinase RNA sequestration

We list the reactions that make up the model for the sRNA-based oscillator, illustrated in Fig. 5B.

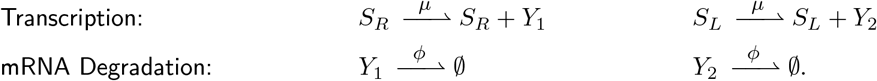

mRNAs *Y*_1_ and *Y*_2_ sequester the recombinases mRNAs *M*_1_ and *M*_2_, respectively, at a rate constant *γ*, so that they become inactive complexes.

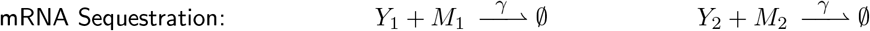

The mRNAs *M*_1_ and *M*_2_ are produced constitutively at a constant rate *θ* and decay with a rate constant *ϕ*. In addition, the mRNAs *M*_1_ and *M*_2_ produce the recombinase monomers *Z*_1_ and *Z*_2_ at a rate constant *ρ*.

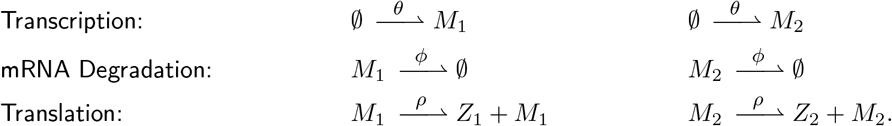

The monomers *Z*_1_ and *Z*_2_ form homodimers *X*_1_ and *X*_2_ as well as the heterodimer, *C*, with an association rate *a* and a dissociation rate *d*. The heterodimer formation is possible because the second recombinase monomer is fused to its RDF. In addition, all protein complexes decay with a rate constant *δ*.

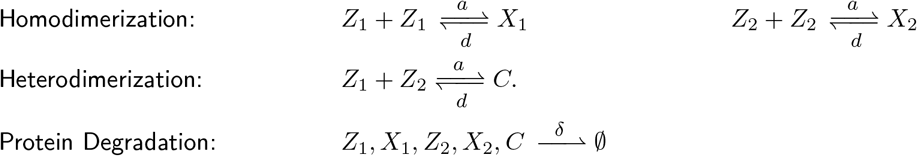

Finally, the inversion rate of the promoter is regulated by the recombinase dimers *X*_1_ and *X*_2_ with rates *f*_1_ and *f*_2_.

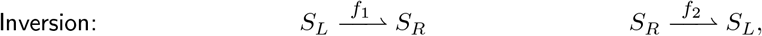

where 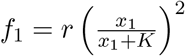 and 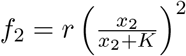.

### Recombinase-based negative feedback oscillator

Fig. 5C shows the design of a negative feedback oscillator using recombinases. It consists of two recombinases, *X*_1_ and *X*_2_, that can invert a promoter. *X*_1_ will invert it to the right and *X*_2_ will invert it back to the left. When the promoter points to the right, only the recombinase *X*_2_ is produced while *X*_1_ is being produced constitutively.

We propose the following reactions as the model of the negative feedback circuit design for the Gille-spie algorithm. The inverting promoter, between recombinase binding sites, regulates the transcription of mRNA *M*_2_, while the constitutive promoter transcribes *M*_1_ at a constant rate of *α*. Both mRNAs decay with a rate constant *ϕ*. In addition, the mRNAs produce recombinase monomers *Z*_1_ and *Z*_2_ with a rate constant *ρ*_1_.

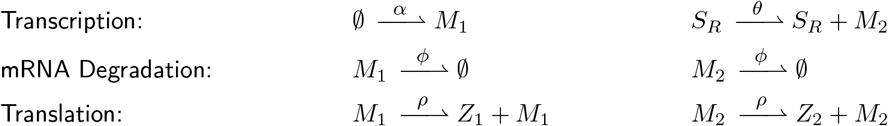

The monomers *Z*_1_ and *Z*_2_ can form homodimers *X*_1_ and *X*_2_ as well as heterodimer, *C*, with an association rate constant *a* and a dissociation rate constant *d*. The heterodimer formation is possible because the second recombinase monomer is fused to the RDF. In addition, all protein complexes decay with a rate constant *δ*.

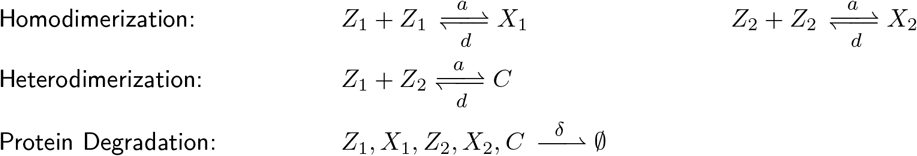

Finally, the rate of promoter inversion is regulated by the recombinase dimers *X*_1_ and *X*_2_ with rates *f*_1_ and *f*_2_.

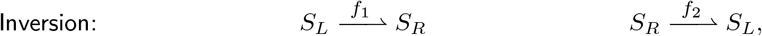

where 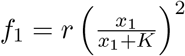 and 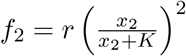.

### Parameter values for simulations

## Acknowledgements

EF and CCS acknowledge support by BBRSC-NSF/BIO award 2020039. JL acknowledges support by the B.I.G. program at UCLA. GG acknowledges support by the Strategic Grant MOSES at the University of Trento.

## References

[1] Kauffman S. Measuring a mitotic oscillator: The ARC discontinuity. Bulletin of Mathematical Biology. 1974;36:171–182.

[2] Goldbeter A. Modeling the mitotic oscillator driving the cell division cycle. Comments on Theoretical Biology. 1993;3:75–107.

[3] Romond PC, Guilmot JM, Goldbeter A. The mitotic oscillator: Temporal self-organization in a phosphorylation-dephosphorylation enzymatic cascade. Berichte der Bunsen-Gesellschaft für Physikalische Chemie. 1994;98:1152–1159.

[4] Barkai N, Leibler S. Circadian clocks limited by noise. Nature. 2000;403(6767):267–268.

[5] Mirsky HP, Liu AC, Welsh DK, Kay SA, Doyle FJ. A model of the cell-autonomous mammalian circadian clock. Proceedings of the National Academy of Sciences. 2009;106(27):11107–11112.

[6] Winfree AT. Human body clocks and the timing of sleep. Nature. 1982;297(5861):23–27.

[7] Uriu K, Morishita Y, Iwasa Y. Traveling wave formation in vertebrate segmentation. Journal of theoretical biology. 2009;257(3):385–396.

[8] Uriu K, Morishita Y, Iwasa Y. Synchronized oscillation of the segmentation clock gene in vertebrate development. Journal of mathematical biology. 2010;61(2):207–229.

[9] Novák B, Tyson JJ. Design principles of biochemical oscillators. Nature reviews Molecular cell biology. 2008;9(12):981–991.

[10] Shitiri E, Vasilakos AV, Cho HS. Biological oscillators in nanonetworksâĂŤOpportunities and challenges. Sensors. 2018;18(5):1544.

[11] Bratsun D, Volfson D, Tsimring LS, Hasty J. Delay-induced stochastic oscillations in gene regulation. Proceedings of the National Academy of Sciences. 2005;102(41):14593–14598.

[12] Gonze D, Abou-Jaoudé W. The Goodwin model: behind the Hill function. PloS one. 2013;8(8):e69573.

[13] Cuba Samaniego C, Giordano G, Kim J, Blanchini F, Franco E. Molecular titration promotes oscillations and bistability in minimal network models with monomeric regulators. ACS synthetic biology. 2016;5(4):321–333.

[14] Blanchini F, Cuba Samaniego C, Franco E, Giordano G. Homogeneous time constants promote oscillations in negative feedback loops. ACS synthetic biology. 2018;7(6):1481–1487.

[15] Purcell O, Savery NJ, Grierson CS, Di Bernardo M. A comparative analysis of synthetic genetic oscillators. Journal of The Royal Society Interface. 2010;7(52):1503–1524.

[16] Potvin-Trottier L, Lord ND, Vinnicombe G, Paulsson J. Synchronous long-term oscillations in a synthetic gene circuit. Nature. 2016;538(7626):514–517.

[17] Riglar DT, Richmond DL, Potvin-Trottier L, Verdegaal AA, Naydich AD, Bakshi S, et al. Bacterial variability in the mammalian gut captured by a single-cell synthetic oscillator. Nature communications. 2019;10(1):1–12.

[18] Santos-Moreno J, Tasiudi E, Stelling J, Schaerli Y. Multistable and dynamic CRISPRi-based synthetic circuits. Nature communications. 2020;11(1):1–8.

[19] Merrick CA, Zhao J, Rosser SJ. Serine integrases: advancing synthetic biology. ACS synthetic biology. 2018;7(2):299–310.

[20] Kim T, Weinberg B, Wong W, Lu TK. Scalable recombinase-based gene expression cascades. Nature Communications. 2021;12(1):1–9.

[21] Guiziou S, Mayonove P, Bonnet J. Hierarchical composition of reliable recombinase logic devices. Nature communications. 2019;10(1):1–7.

[22] Pokhilko A, Ebenhöh O, Stark WM, Colloms SD. Mathematical model of a serine integrase-controlled toggle switch with a single input. Journal of The Royal Society Interface. 2018;15(143):20180160.

[23] Zhao J, Pokhilko A, Ebenhöh O, Rosser SJ, Colloms SD. A single-input binary counting module based on serine integrase site-specific recombination. Nucleic acids research. 2019;47(9):4896–4909.

[24] Samaniego CC, Giordano G, Franco E. Periodic switching in a recombinase-based molecular circuit. IEEE Control Systems Letters. 2019;4(1):241–246.

[25] Meinke G, Bohm A, Hauber J, Pisabarro MT, Buchholz F. Cre recombinase and other tyrosine recombinases. Chemical reviews. 2016;116(20):12785–12820.

[26] Siuti P, Yazbek J, Lu TK. Synthetic circuits integrating logic and memory in living cells. Nature biotechnology. 2013;31(5):448–452.

[27] Bonnet J, Subsoontorn P, Endy D. Rewritable digital data storage in live cells via engineered control of recombination directionality. Proceedings of the National Academy of Sciences. 2012;109(23):8884–8889.

[28] Fernandez-Rodriguez J, Yang L, Gorochowski TE, Gordon DB, Voigt CA. Memory and combinatorial logic based on DNA inversions: dynamics and evolutionary stability. ACS synthetic biology. 2015;4(12):1361–1372.

[29] Folliard T, Steel H, Prescott TP, Wadhams G, Rothschild LJ, Papachristodoulou A. A synthetic recombinase-based feedback loop results in robust expression. ACS synthetic biology. 2017;6(9):1663–1671.

[30] Steel H, Papachristodoulou A. Low-burden biological feedback controllers for near-perfect adaptation. ACS synthetic biology. 2019;8(10):2212–2219.

[31] Weinberg BH, Pham NH, Caraballo LD, Lozanoski T, Engel A, Bhatia S, et al. Large-scale design of robust genetic circuits with multiple inputs and outputs for mammalian cells. Nature biotechnology. 2017;35(5):453–462.

[32] Weinberg BH, Cho JH, Agarwal Y, Pham NH, Caraballo LD, Walkosz M, et al. High-performance chemical-and light-inducible recombinases in mammalian cells and mice. Nature communications. 2019;10(1):1–10.

[33] Gillespie DT. Exact stochastic simulation of coupled chemical reactions. The journal of physical chemistry. 1977;81(25):2340–2361.

[34] Kuo J, Yuan R, Sánchez C, Paulsson J, Silver PA. Toward a translationally independent RNA-based synthetic oscillator using deactivated CRISPR-Cas. Nucleic acids research. 2020;48(14):8165–8177.

[35] Yan J. Identifying Hard Bounds on Molecular Fluctuation in Stochastic Reaction Systems. Harvard University; 2020.

[36] Olorunniji FJ, McPherson AL, Rosser SJ, Smith MC, Colloms SD, Stark WM. Control of serine integrase recombination directionality by fusion with the directionality factor. Nucleic acids research. 2017;45(14):8635–8645.

[37] Glass DS, Jin X, Riedel-Kruse IH. Nonlinear delay differential equations and their application to modeling biological network motifs. Nature communications. 2021;12(1):1–19.

[38] Gao XJ, Chong LS, Kim MS, Elowitz MB. Programmable protein circuits in living cells. Science. 2018;361(6408):1252–1258.

[39] Fink T, Lonzarić J, Praznik A, Plaper T, Merljak E, Leben K, et al. Design of fast proteolysis-based signaling and logic circuits in mammalian cells. Nature chemical biology. 2019;15(2):115–122.

[40] Fern J, Scalise D, Cangialosi A, Howie D, Potters L, Schulman R. DNA strand-displacement timer circuits. ACS synthetic biology. 2017;6(2):190–193.

[41] Schlosshauer M, Baker D. Realistic protein–protein association rates from a simple diffusional model neglecting long-range interactions, free energy barriers, and landscape ruggedness. Protein science. 2004;13(6):1660–1669.

[42] Schreiber G, Haran G, Zhou HX. Fundamental aspects of protein-protein association kinetics. Chemical reviews. 2009;109(3):839–860.

[43] Buchler NE, Louis M. Molecular titration and ultrasensitivity in regulatory networks. Journal of molecular biology. 2008;384(5):1106–1119.

[44] Kim J, Khetarpal I, Sen S, Murray RM. Synthetic circuit for exact adaptation and fold-change detection. Nucleic acids research. 2014;42(9):6078–6089.

[45] Andersen JB, Sternberg C, Poulsen LK, Bjørn SP, Givskov M, Molin S. New unstable variants of green fluorescent protein for studies of transient gene expression in bacteria. Applied and environmental microbiology. 1998;64(6):2240–2246.

[46] Proshkin S, Rahmouni AR, Mironov A, Nudler E. Cooperation between translating ribosomes and RNA polymerase in transcription elongation. Science. 2010;328(5977):504–508.

[47] Basu S, Gerchman Y, Collins CH, Arnold FH, Weiss R. A synthetic multicellular system for programmed pattern formation. Nature. 2005;434(7037):1130–1134.

[48] Qian Y, Huang HH, Jiménez JI, Del Vecchio D. Resource competition shapes the response of genetic circuits. ACS synthetic biology. 2017;6(7):1263–1272.

[49] Liu M, Gupte G, Roy S, Bandwar RP, Patel SS, Garges S. Kinetics of transcription initiation at lacP1: Multiple roles of cyclic AMP receptor protein. Journal of Biological Chemistry. 2003;278(41):39755–39761.

[50] Milo R, Jorgensen P, Moran U, Weber G, Springer M. BioNumbersâĂŤthe database of key numbers in molecular and cell biology. Nucleic acids research. 2010;38(suppl_1):D750–D753.

[51] Pokhilko A, Zhao J, Ebenhöh O, Smith MC, Stark WM, Colloms SD. The mechanism of *ϕ*C31 integrase directionality: experimental analysis and computational modelling. Nucleic acids research. 2016;44(15):7360–7372.

[52] Yi L, Gebhard MC, Li Q, Taft JM, Georgiou G, Iverson BL. Engineering of TEV protease variants by yeast ER sequestration screening (YESS) of combinatorial libraries. Proceedings of the National Academy of Sciences. 2013;110(18):7229–7234.

[53] Kim J, White KS, Winfree E. Construction of an in vitro bistable circuit from synthetic transcriptional switches. Molecular systems biology. 2006;2(1):68.

[54] Zhang DY, Turberfield AJ, Yurke B, Winfree E. Engineering entropy-driven reactions and networks catalyzed by DNA. Science. 2007;318(5853):1121–1125.

